# Inkject Printed Graphene Microelectrodes for Real-Time Impedance Spectroscopy of Neural Cells in Organ-on-a-Chip

**DOI:** 10.1101/2024.02.28.582633

**Authors:** Lionel Jean Gabriel Ouedraogo, Nicole N. Hashemi

**Affiliations:** Department of Mechanical Engineering, Iowa State University, Ames, IA, 50011, USA; Department of Biomedical Sciences, Iowa State University, Ames, IA, 50011, USA

**Keywords:** organ-on-a-chip, brain-on-a-chip, in vitro, microfluidics

## Abstract

This study presents the development and characterization of a graphene-based sensor integrated into a microfluidic chip for real-time monitoring of cell growth and viability. The sensor fabrication involved the metabolization of graphene from graphite using a simple and cost-effective method. The sensor design, created using Solidworks, featured electrodes capable of detecting environmental changes through impedance sensing. A mold was created using a cutter plotter to overcome challenges in achieving the desired sensor shape, and the electrodes were printed on a polyester (PETE) membrane. The conductivity of the electrodes was optimized through annealing, considering the temperature limits of the membrane. Annealing at 150 °C for 40 minutes yielded electrodes with desired conductivity and maintained membrane integrity. Compatibility with cellular growth was confirmed through cell culture experiments. The scaled electrodes were integrated into a microfluidic chip, and their performance was evaluated using cyclic voltammetry and electrochemical impedance spectroscopy. The results demonstrated the successful functioning of the electrodes within the chip. The developed graphene-based sensor offers promising applications in cellular studies and biosensing through real-time monitoring of cell growth and viability was achieved by measuring impedance changes resulting from cell attachment.

## 1. Introduction

Graphene, a two-dimensional material composed of a single layer of carbon atoms arranged in a hexagonal lattice, has emerged as a revolutionary material with exceptional properties [1]. Its unique combination of mechanical strength, electrical conductivity, and thermal stability has made it a subject of intense research across various scientific and technological fields. One area where graphene has shown tremendous potential is in the development of organ-on-a-chip (OOC) systems [2,3], which aim to mimic the structure and function of human organs in a laboratory setting. The integration of sensors into these OOC systems is crucial for enhancing data-gathering capabilities, leading to a better understanding of organ behavior and response. In this study, we focus on the utilization of pristine graphene to fabricate sensors for integration into an organ-on-a-chip platform. Graphene-based sensors offer numerous advantages, including high electrical conductivity, biocompatibility, and sensitivity [4], making them an ideal choice for real-time monitoring and analysis of organ behavior. By incorporating graphene-based sensors into OOC platforms, researchers can obtain continuous, accurate, and non-invasive data on various physiological parameters, such as temperature [5], pH [6], oxygen levels [7,8], and specific biomarkers [9,10,11].

The development of these graphene-based sensors involves several key steps, including the synthesis of graphene and the design and fabrication of the sensors themselves. In our study, we synthesized graphene through a simple and cost-effective process, metabolizing it from graphite. A solution containing graphite and bovine serum albumin (BSA) was prepared, followed by ball milling and sedimentation to remove impurities. The quality of the resultant graphene was characterized using Raman spectroscopy and scanning electron microscopy (SEM).

To fabricate the sensors, a design was created using Solidworks software, incorporating a simple yet effective layout capable of detecting environmental changes through impedance sensing. The electrodes were printed onto a substrate using an inkjet printer, overcoming initial challenges by employing a mold that facilitated the production of the desired sensor configuration. Post-processing techniques, such as annealing, were employed to enhance the conductivity of the electrodes while ensuring the compatibility with the membrane material used in the OOC system. The optimal annealing conditions were determined by evaluating the resistance of the electrodes at various temperatures and durations.

Integration of the graphene-based sensors into the OOC system involved the fabrication of a microfluidic chip, which serves as the platform for culturing and studying organ behavior. The chip was constructed using a SU-8 mold and PDMS layers, with a porous membrane sandwiched between them. The graphene electrodes were positioned on the membrane, and the layers were bonded together to create a leak-free chip.

The successful integration of the pristine graphene sensor within the OOC platform enabled real-time monitoring of organ behavior and response. The sensors’ electrochemical performance was analyzed using techniques such as cyclic voltammetry and electrochemical impedance spectroscopy, which provided insights into their stability, sensitivity, and functionality.

The successful integration and performance of the graphene-based sensors within the OOC system open up exciting opportunities for advancing research in various fields, including organ physiology, drug testing, disease modeling, and personalized medicine.

## 2. Experimental Section

### 2.1 Graphene synthesis

During the study, graphene was metabolized from graphite. The process employed was characterized by its simplicity and cost-effectiveness [12]. A solution was prepared by combining 20 mg/mL graphite powder (particle size <20 μm, synthetic, Sigma-Aldrich) and 2 mg/mL BSA (lyophilized powder, Sigma-Aldrich) in deionized water. The solution was subjected to ball milling at a rotational speed of 300 rpm for 90 hours using stainless-steel balls (11/32″). Afterward, the solution was allowed to rest for 48 hours to allow sedimentation of large graphite particles. To remove Fe impurities, 85% of the solution volume from the top was pipetted off. Raman spectra **figure 1 a)** were obtained using a confocal Raman system (BWTEK Voyage) with a CW laser (Excelsior-532-150-CDRH Spectra-Physics) operating at a wavelength of 532 nm to characterize the produced graphene. The aqueous graphene was drop-casted onto Si/SiO2 wafers with a drop diameter of approximately 10 mm. Raman data were collected from five different laser spots across five independently printed electrodes and five untreated grapheme samples (n = 30) with a laser integration time of 10 seconds. Scanning electron microscope (SEM) images of the printed graphene patterns on PETE were obtained using a JEOL FESM 6335 at 2–5 kV accelerating voltage **figure 2**. The impedance and cyclic voltammetry measurements were carried out using a VersaSTAT 4 Potentiostat Galvanostat **figure 4**.

**Figure 1.**
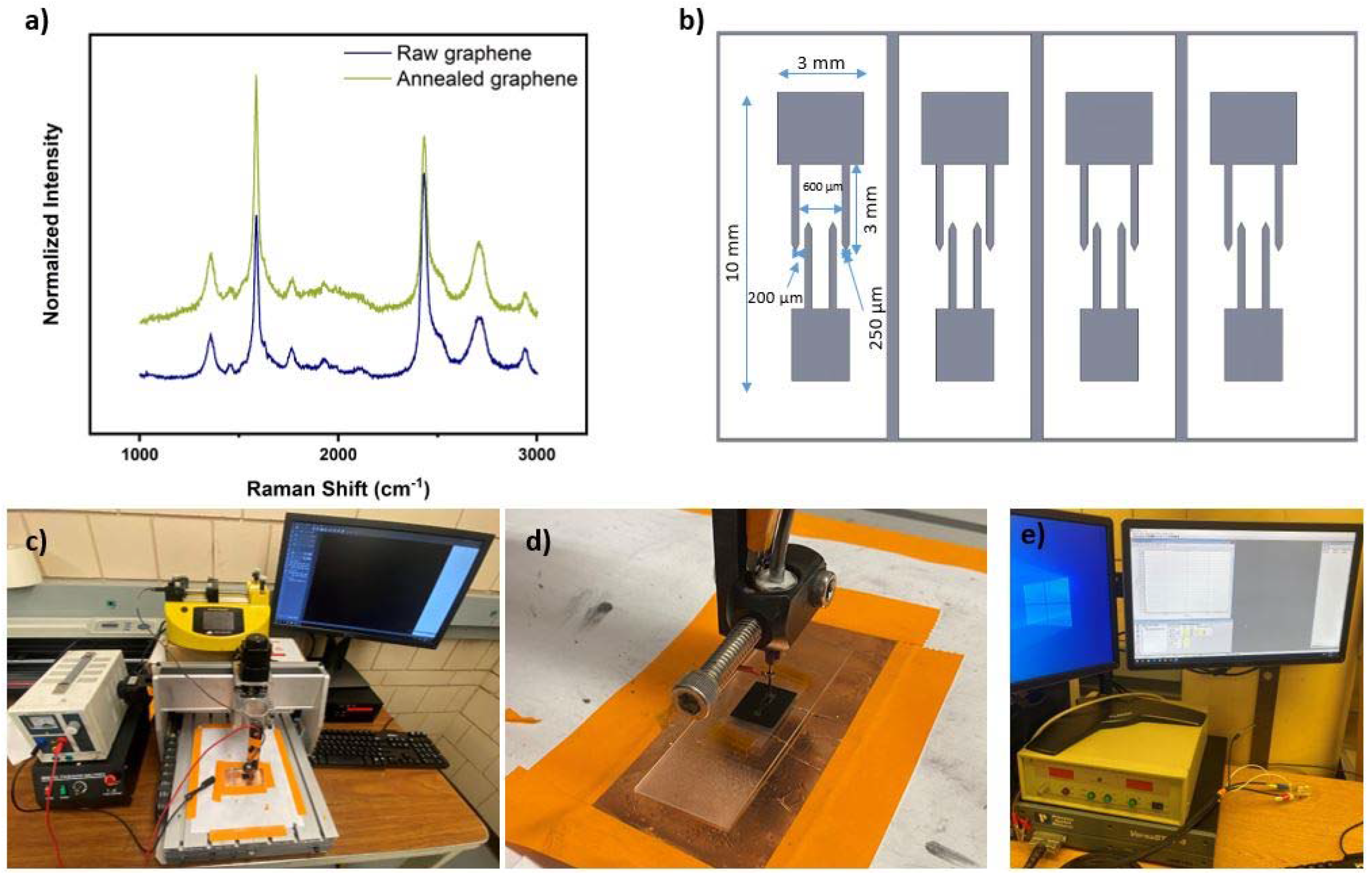
**a)** Raman shifts of pretreated and thermally annealed samples acquired at 532 nm. **b)** schematics and dimensional characteristics pertaining to the interdigitated electrode (IDE) structure. **c)** The experimental setup involved the utilization of a syringe pump in conjunction with an inkjet printer for controlled ink deposition. The ink was deposited at a constant flow rate of 9 μL/s to ensure uniformity and consistency in the printing process. **d)** The needle and substrate setup, a potential difference of 3 kV was applied between the substrate and the needle to facilitate the deposition of ink onto the substrate surface. To ensure stability and support, a cover glass was positioned on a conventional microscope slide. This arrangement provided a suitable platform for the ink affixation process. **e)** Impedance Spectroscopy and Cyclic Voltammetry setup with Biosensor.

**Figure 2.**
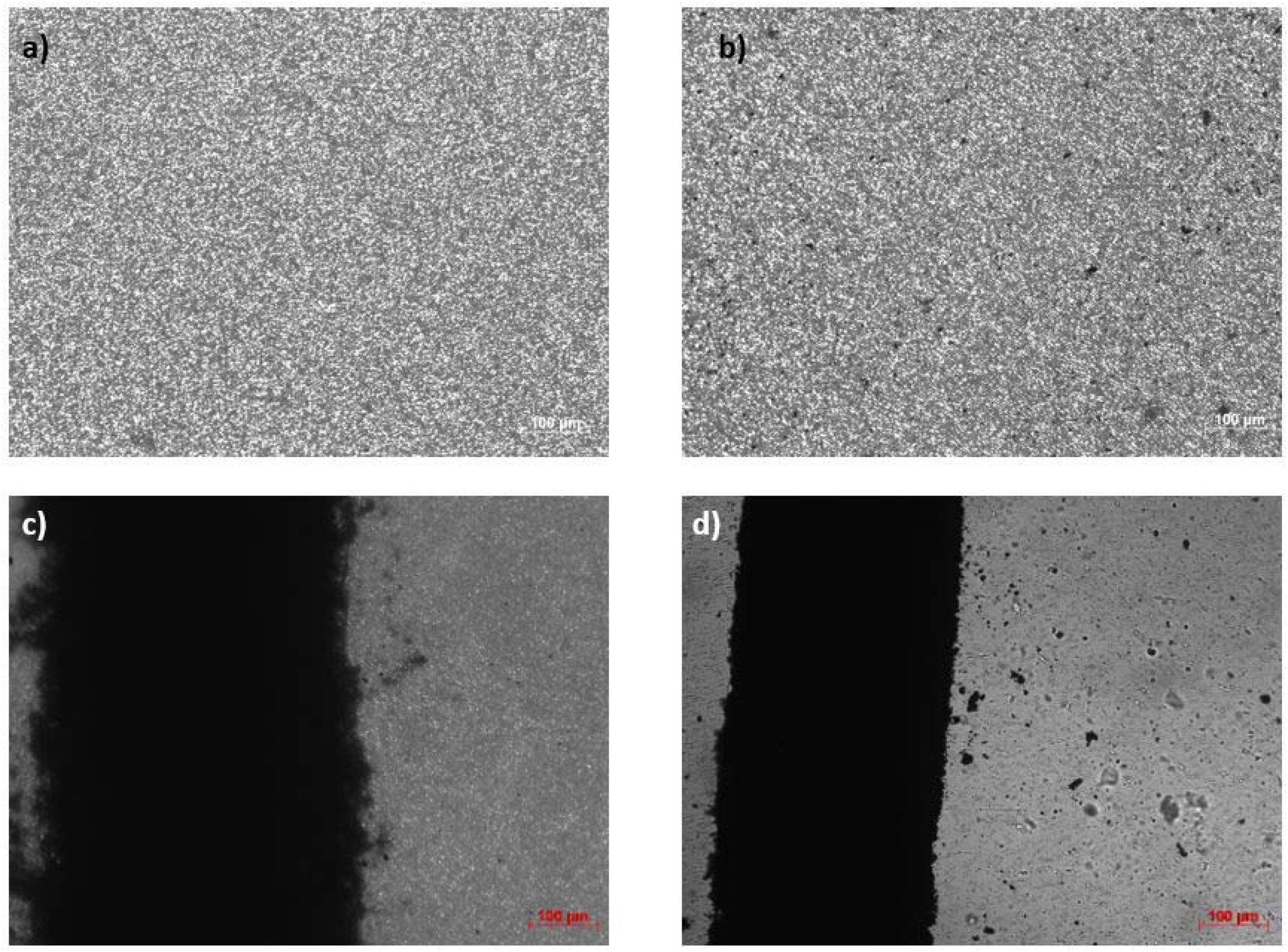
**a)** Microscopy imaging of membrane before annealing **b)** Microscopy imaging of membrane after annealing at 150 degrees Celsius for 40 minutes **c)** Microscopy imaging of graphene printed on opaque membrane. **d)** Microscopy imaging of graphene printed on transparent membrane.

**Figure 3.**
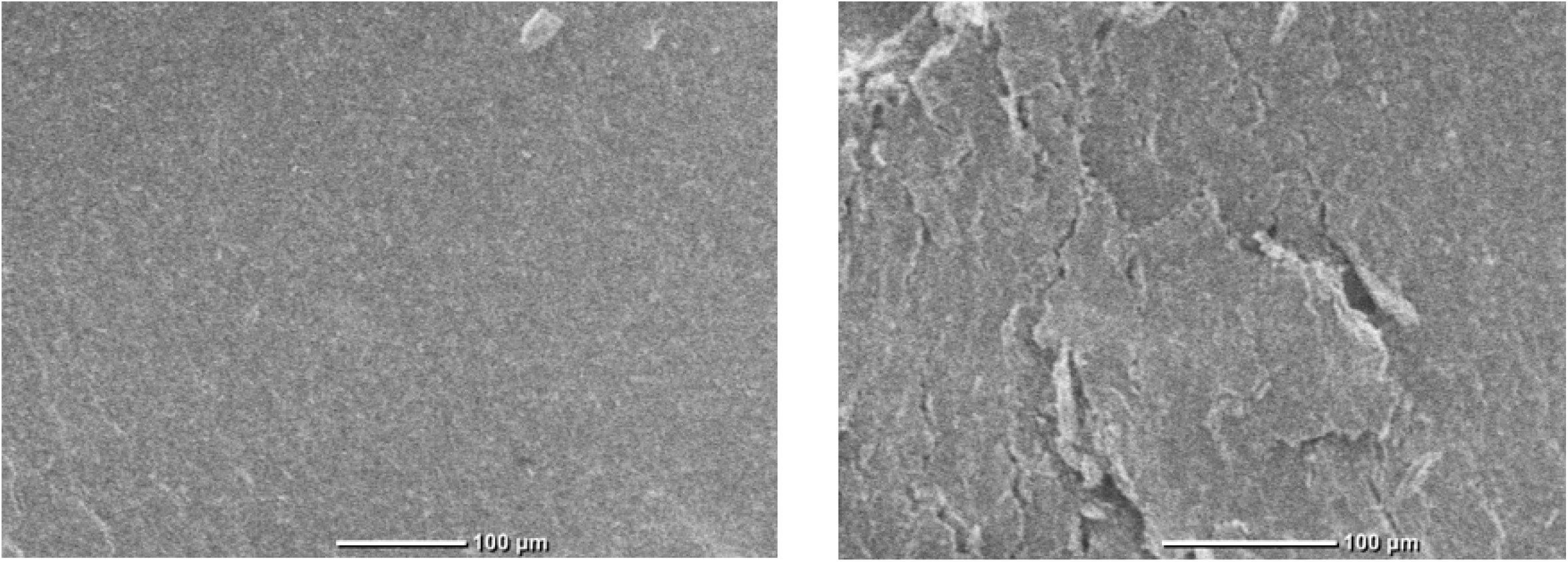
**a)** SEM imaging of graphene electrode before annealing **b)** SEM image of graphene electrodes after annealed for 60 minutes at 150 degrees Celsius.

**Figure 4.**
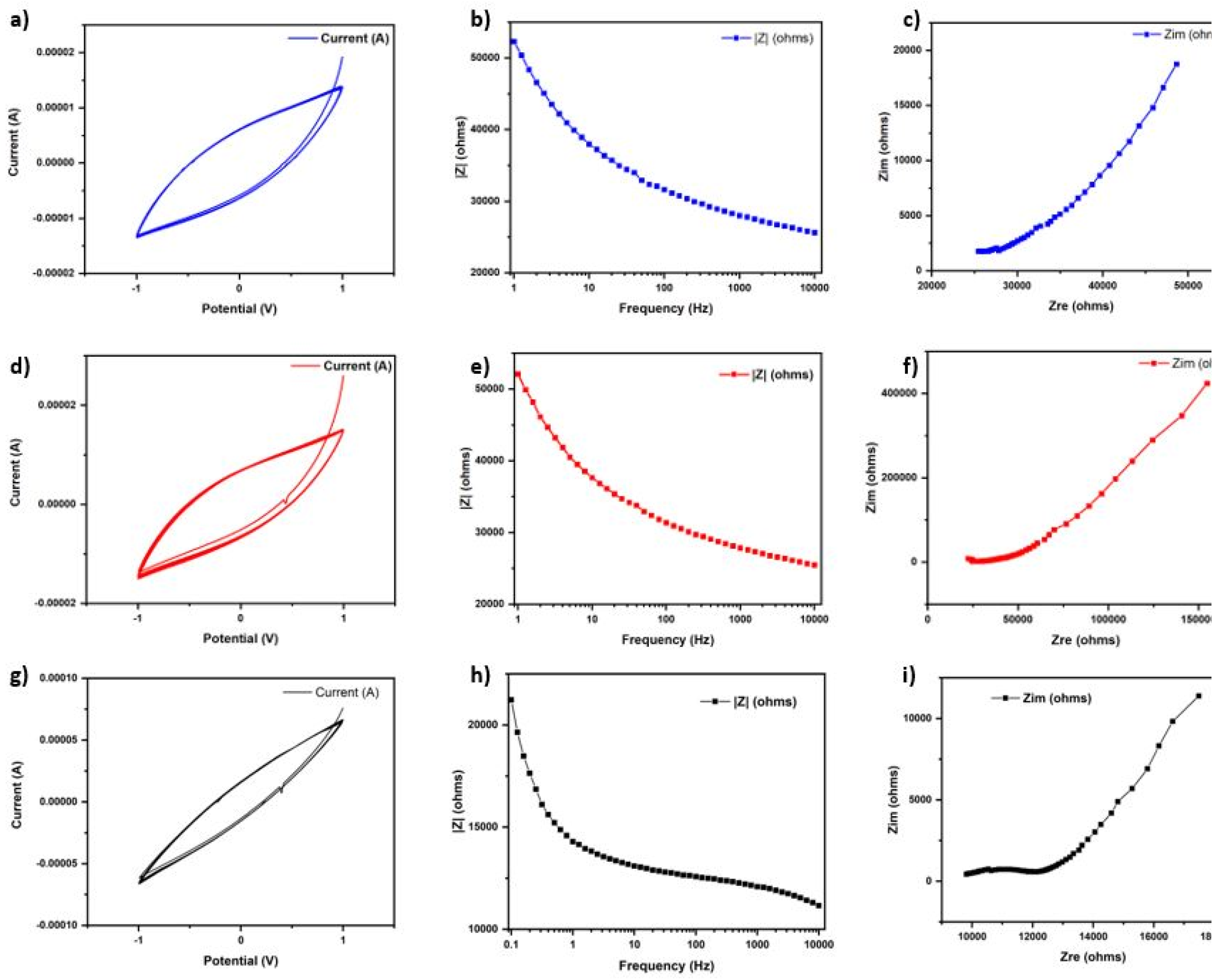
Characterization comparison between macro and micro electrodes. **a)** Cyclic voltammetry of macro electrodes in KCL. **b)** Impedance spectroscopy of macro electrodes in KCL. **c)** Nyquist plot, Z_re_ vs Z_im_ of macro electrodes. **d)** Cyclic voltammetry of micro electrodes in KCL. **e)** Impedance spectroscopy of micro electrodes in KCL. **f)** Nyquist plot, Z_re_ vs Z_im_ of micro electrodes. **g)** Cyclic voltammetry of micro electrodes in MM. **h)** Impedance spectroscopy of micro electrodes in MM. **i)** Nyquist plot, Z_re_ vs Z_im_ of micro electrodes.

### 2.2 Design and Fabrication of the Macroelectrodes

Interdigitated electrodes were designed utilizing Solidworks software, incorporating a simple yet effective design capable of detecting environmental changes through impedance sensing **figure 1 b)**. Interdigitated electrodes exhibit a high surface area-to-volume ratio, allowing for improved sensitivity in detecting and measuring the signals and analytes. The closely spaced finger-like structures create a larger electrode-electrolyte interface, facilitating efficient electrochemical reactions and enhancing signal detection. The interdigitated electrode design minimizes the capacitive coupling between adjacent fingers, reducing the interference and noise caused by stray capacitance. This feature improves the signal-to-noise ratio and enables accurate measurements of weak electrical signals. The closely spaced fingers of interdigitated electrodes promote efficient charge transfer and reduce the diffusion distance for ions or molecules in the electrolyte solution. This design enhances the electrode’s electrochemical performance, allowing for faster and more efficient reactions.

To fabricate the electrodes, a method was devised involving the use of an inkjet printer. Interdigitated electrodes can be fabricated in micro or nano-scale dimensions, enabling miniaturization of devices and integration into compact systems. This feature is particularly advantageous for our applications where space is limited. However, the process encountered challenges initially when attempting to directly deposit the sensors onto the scaffold, resulting in difficulties achieving the desired shape. To overcome this hurdle, we opted to create a mold. The dimensions of the sensors were extracted from Solidworks and utilized to precisely cut the mold using a US cutter plotter. Once the mold was obtained, it was positioned on the scaffold, enabling successful printing of the electrodes. Although the process involved a significant duration, the utilization of the mold ultimately facilitated the production of the desired sensor configuration. The experimental setup involved the utilization of a syringe pump in conjunction with an inkjet printer for controlled ink deposition **Figure 1 c)**. The ink was deposited at a constant flow rate of 9 μL/s to ensure uniformity and consistency in the printing process. A potential difference of 3 kV was applied between the substrate and the needle to facilitate the deposition of ink onto the substrate surface **figure 1 d)**. To ensure stability and support, a cover glass was positioned on a conventional microscope slide. This arrangement provided a suitable platform for the ink affixation process.

### 2.3 Post-Processing

We identified that the conductivity of the electrodes did not meet desired specifications. To enhance the electrode conductivity, we implemented an annealing process. However, due to the utilization of a polyester membrane for printing, we needed to ensure that the annealing process did not exceed certain temperatures that could adversely affect the membrane properties. To determine the safe limit for annealing, we conducted incremental temperature testing, ranging from 110 to 200 degrees Celsius. Notably, we observed an increase in the stiffness of the material at 160 degrees Celsius. Consequently, we decided to set the annealing temperature for the graphene electrodes at 150 degrees Celsius.

Furthermore, we explored the impact of annealing time on the membrane’s properties. We subjected the material to temperatures of 150 degrees Celsius for varying durations, ranging from 10 to 60 minutes with 10 minutes’ increments. Through stiffness testing, we found that the material’s stiffness remained relatively consistent after 60 minutes of heating at 150 degrees Celsius. We observed two membranes under the microscope to assess changes in porosity. One that had been annealed at 150 degrees Celsius for 60 minutes’ **figure 2 a)** and another that had not undergone annealing **figure 2 b)**.

To assess the effect of annealing on cellular growth, we conducted a cell culture experiment. We cultured cells on two membranes: one that had been annealed at 150 degrees Celsius for 60 minutes and another that had not undergone annealing. After monitoring the cell culture for three days, we observed that the cells exhibited similar growth rates on both membranes, indicating that annealing the membrane at the designated temperature did not impact cellular growth.

Once we determined the limit temperature for annealing, we proceeded to conduct annealing tests on the graphene electrodes. The annealing temperatures ranged from 100 degrees to 150 degrees Celsius. Through observations, we noted that the resistance of the electrodes decreased as the temperature increased. The lowest resistance was achieved at 150 degrees Celsius, indicating the optimal annealing temperature.

Recognizing the influence of annealing time on resistance, we further investigated its impact. We annealed the electrodes at 150 degrees Celsius for durations ranging from 10 to 60 minutes in 10-minute increments. It was observed that the resistance decreased over time, reaching its lowest point after 40 minutes of annealing. However, beyond 40 minutes, the resistance began to increase. To gain insights into this phenomenon, we conducted additional analyses using a scanning electron microscope (SEM). Upon observation, we discovered no crack on graphene annealed for 40 minutes’ **figure 3 a)** and the presence of cracks on the surface of the electrodes annealed for 60 minutes’ **figure 3 b)**.

Taking all these findings into account, we determined the optimal annealing conditions for the graphene electrodes to be 150 degrees Celsius for 40 minutes. Under these conditions, the electrodes exhibited a resistance of approximately 51 Ohms sqr-1 while maintaining the integrity of the membrane.

### 2.4 Rescaling

Using Solidworks, the sensor redesign to align with the dimensions of the organ-on-a-chip. The design was modified to overlap the chip’s channel, enabling impedance measurements within the channel. Additionally, the electrode extended beyond the channel, allowing PDMS to act as a passivating medium. The electrode ends were positioned at a midpoint between the channel and the chip’s edge to ensure secure bonding and prevent leaks. Ample space was left to create access holes for the copper wires.

To fabricate the new electrodes, molds were created using a cutter plotter. However, challenges arose during the initial attempts as the cutter plotter encountered difficulties in cutting the vinyl to the required shape due to the small dimensions of the microelectrodes. Adjustments were made to the parameters and experiment with different settings to achieve the desired results. Once the optimal parameters were determined, molds were consistently produced without any issues. Subsequently, the electrodes were printed using the molds.

By overcoming the challenges and successfully fabricating the new electrodes, we made significant progress towards integrating the scaled electrodes into the microfluidic chip. This advancement would enable precise impedance measurements within the chip’s channel, contributing to the further development and functionality of the organ-on-a-chip system. Both macro and micro electrodes were characterized by CV, EIS and Nyquist plot and showed very similar properties and capacities **figure 4**.

### 2.5 Chip fabrication and sensor integration

To construct the microfluidic chip, a SU-8 mold was created using soft lithography, allowing for the fabrication of the top and bottom layers with channels. The chip comprises a top and bottom layer with a porous PETE membrane (2×10 mm) sandwiched between them [13-15]. Graphene electrodes were printed on top of the membrane prior to assembly.

The layers of the chip were formed by pouring a PDMS solution mixed with a curing agent (in a 10:1 ratio) into the mold. The solution was then left to cure for 48 hours. Once cured, the layers were carefully removed from the mold, and the membrane with the printed electrodes was positioned at the center of the chip.

Inlet and outlet holes were created on the top layer using a 1 mm biopsy punch. On the bottom layer, two holes were made using a 1.5 mm biopsy punch to allow for the connection of copper wires with the electrodes. Initially, the wires were laid between the top and bottom layers, but this method resulted in leaks at the interfaces. It was found that clean and perfectly flat surfaces were necessary for successful bonding. To address this issue, the copper wires were fed through the access holes on the bottom layer and connected to the electrodes using silver paste.

The top and bottom layers were bonded together using a PDMS-to-PDMS bonding technique. This involved cleaning the bonding surfaces with pressurized air to remove debris and treating the surfaces in a Plasma cleaner to activate them. The layers were then pressed together and left to bond overnight. Once the bonding was complete, the access holes on the bottom layer were sealed with epoxy to prevent leaks, contamination, and potential junction fractures during operation.

For fluid access, 1/16 ft PEEK tubes were inserted into each inlet and outlet hole on the top layer. Finally, a leak test was conducted using a solution of DI water and green dye to ensure the chip’s integrity and to verify that there were no leaks in the system.

This comprehensive assembly process ensured the successful integration of the printed graphene electrodes within the microfluidic chip, providing a functional platform for further experiments and analysis.

### 2.6 Testing the sensors in the microfluidic chip

To conduct cyclic voltammetry (CV), a potentiostat was used in conjunction with the microfluidic chip. A silver electrode was employed as the reference electrode, while the graphene electrodes within the chip served as both the working and counter electrodes. As the electrolyte, a potassium chloride (KCl) solution was prepared by mixing deionized (DI) water with one mole of KCl.

The KCl solution was introduced into the channels of the microfluidic chip, and CV measurements were performed within a potential range of 1 to 1 Volt, using a sampling rate of 10 millivolts per second. This allowed for the observation of faradaic peaks, which are indicative of electrochemical reactions occurring at the electrodes **figure 4 d)**.

Furthermore, electrochemical impedance spectroscopy (EIS) was conducted using the same experimental setup. The measurements were carried out at various intervals, ranging from 0.1 Hz to 10,000 Hz, while maintaining a direct current (Vdc) of 0 mV and an alternating current (Vac) of 10 mV. EIS provides information about the impedance of the electrodes at different frequencies, allowing for the characterization of their electrical properties **figure 4 e)**.

Based on the results obtained from these electrochemical tests, it was determined that the graphene electrodes within the microfluidic chip were functioning as intended. This indicates that the electrodes were successfully integrated into the chip and capable of detecting changes in impedance, providing a foundation for further analysis and experimentation.

## 3. Results and Discussion

### 3.1 Biological and physical characterization

Before conducting neural studies using our membrane with deposited graphene, we had to evaluate the physical, biological, and chemical properties of the membrane. In a previous study, our sensor demonstrated improved stability and conductivity after post-treatment. The optimal annealing temperature for post-processing was determined to be 280 °C for 30 minutes. However, since the membrane used as a scaffold in this study couldn’t withstand such high temperatures without compromising its integrity, we modified the protocol to suit our requirements.

We printed graphene on PETE and annealed it at temperatures ranging from 100 °C to 150 °C for 30 minutes. The resistance of the graphene decreased as the annealing temperature increased. After assessing various temperatures, we concluded that annealing the deposited graphene at 150 °C was ideal as it enhanced electrode conductivity while preserving the membrane’s integrity and removing impurities. To investigate the effect of annealing time on resistance, we printed five graphene lines on PETE and annealed them for different durations (30, 35, 40, 45, and 50 minutes) at 150 °C. The average resistance decreased until 40 minutes, but further annealing caused a reduction in conductivity.

To understand why the conductivity decreased after 40 minutes of annealing, we conducted further investigation. SEM analysis **Figure 3** revealed cracks and fissures in the electrodes annealed for 60 minutes, which negatively impacted conductivity. The electrical conductivity of thin films is affected by surface roughness, resulting in increased electric resistance [16]. Annealing the material at 150 °C for 40 minutes did not alter the material properties significantly, including porosity, flexibility, and elasticity of the PETE membrane.

Subsequently, we needed to ensure that the heat treatment did not affect any properties that could influence cellular growth on the chip. The membrane was treated with ECL to enhance cellular attachment. The modified membrane was placed in a 5 cm diameter petri dish, and mouse astrocytes were grown on it for five days using established protocols for mouse astrocyte subculture. Through microscopic observation, we noted that the cells achieved full confluence in the dish. Interestingly, the cells tended to grow around the graphene first before spreading throughout the dish. The cells in proximity to the graphene electrodes exhibited delayed detachment and were the last to die off.

### 3.2 Electrochemical performance analysis

To assess the stability, sensitivity, and electrochemical behavior of the membrane-bound integrated impedance, we conducted spectroscopy and cyclic voltammetry on the macro and micro sensors. Cyclic voltammetry was performed in a 1 M KCl solution with results shown in **figure 4 a)** and **d)** with a scan rate of 100 mV/s, ranging from -1 V to 1 V. Cyclic voltammetry was also performed with MM with results shown in f**igure 4 g)** with a scan rate of 100 mV/s, ranging from -1 V to 1 V. The results, shown in **figure 4 a) d)** and **g)**, exhibited a reversible and rapid transfer of electrons, indicating excellent stability across eighteen cycles. Notably, distinct peaks in the cyclic voltammogram corresponded to the oxidation and reduction of the analyte, demonstrating the efficient electron transfer during the redox process. The well-defined and symmetrical peaks suggested a reversible reaction. Impedance spectroscopy was performed in 1 M KCl in **figure 4 b)** and **e)** and in MM in **figure 4 h)**. The impedance spectroscopy showed the capacity of both macroelectrodes and microelectrodes to detect analytes in KCL and MM and show the growth of cell in MM by sensing the impedance change in the OOC channels. Nyquist plots were obtained with 1 M KCl solution in **figure 4 c)** and **f)** and in MM in **figure 4 i)**. The Nyquist plots in **figure 4 c) f)** and **i)** revealed super capacitors capable of rapid charge\discharge cycles. Supercapacitors exhibit high power density, enabling them to charge and discharge rapidly. This feature is particularly advantageous for sensing applications where real-time monitoring is essential. The rapid charging and discharging of a supercapacitor facilitate quick response times, allowing for precise and timely measurements of cell growth dynamics.

## 4. Conclusion

This study successfully developed and characterized a graphene-based sensor integrated into a microfluidic chip for real-time monitoring of cell growth. The fabrication process involved metabolizing graphene from graphite, and the sensor design was created using Solidworks, enabling the detection of environmental changes through impedance sensing. The conductivity of the electrodes was optimized through annealing, considering the temperature limits of the membrane.

The scaled down microelectrodes were successfully integrated into a microfluidic chip, and their performance was evaluated using cyclic voltammetry and electrochemical impedance spectroscopy. The results indicated the reliable functioning of the electrodes within the chip, providing a base for real-time monitoring of cell growth and viability by measuring impedance changes resulting from cell attachment and membrane integrity.

The developed graphene-based sensor holds great promise in various applications, particularly in cellular studies and biosensing. Its ability to monitor cell growth and viability in real-time offers valuable insights for biological research, drug discovery, and tissue engineering. The integration of the sensor into a microfluidic chip enhances its practicality and opens doors to more sophisticated and high-throughput experiments.

Overall, the developed graphene-based sensor integrated into a microfluidic chip demonstrates its potential as a powerful tool for real-time monitoring of cell growth and viability, contributing to advancements in biomedical research and applications.

## Data Availability

The data that support the findings of this study are available upon reasonable request from the authors.

## Acknowledgments

This work was partially supported by the National Science Foundation Award 2321975 and 2014346.

